# Do plasmid dependent phages promote the survival of costly plasmids?

**DOI:** 10.64898/2026.06.01.729315

**Authors:** Jonathon L. Yuly, Matthew Avallone, Lawrence Abad, Ned S. Wingreen

## Abstract

Plasmids store auxiliary genes that address environmental challenges such as antibiotics. In the short term, plasmid loss can be advantageous to hosts if plasmid benefits are temporary but costs are permanent. However, this loss can proceed to plasmid extinction, with access to plasmid-derived benefits permanently lost. We consider a worst-case scenario: a plasmid that yields benefits only infrequently, imposes heavy costs that cannot be alleviated, and whose host population faces a competitor that cannot acquire the costly plasmid. Thus, horizontal transmission of this plasmid is detrimental to the host population when the plasmid does not provide benefits. Using population models, we demonstrate that – counterintuitively – plasmid-dependent phages can enable high-cost plasmid survival. Phage predation pins the plasmid at low abundance, effectively neutralizing the plasmid cost at the population level and boosting persistence time. When the costly plasmid becomes beneficial, it spreads and survives by vertical transmission until benefits subside.

**Importance Statement:** Conjugative plasmids are often viewed as a mobile “gene library” that can spread beneficial traits to address environmental challenges. When a bacterial community encounters environmental challenges such as antibiotics, plasmids at low abundance can quickly spread and rescue the population. But what prevents the extinction of these plasmids between selection events, when they convey costs but not benefits? We propose that plasmid-dependent phages can – counterintuitively – promote the survival of high-cost plasmids. By targeting conjugative machinery, these phages suppress costly plasmids to low but nonzero abundance, dramatically reducing the effective cost of the plasmid to the host population. Under realistic levels of phage infectivity, predation also becomes self-limiting as plasmid-bearing hosts become scarce. Together these factors can enable otherwise high-cost plasmids to persist indefinitely.

## Main text

Bacteria in nature face fierce competition across diverse and often rapidly changing environments. To help them survive, bacteria harbor and exchange plasmids, small genetic elements that can encode auxiliary genes to address transient or niche environmental challenges [1, 2, 3, 4]. For example, if a community is challenged by an antibiotic or toxin, plasmids with genes that provide resistance can spread and rescue the population. Similarly, if a niche energy or nutrient source is encountered, plasmids can spread that encode alternate metabolic pathways [5, 6, 7]. Should these environmental challenges subside, the population can lose the plasmid and any associated costs without relying on chromosomal modifications. However, positive selection for a plasmid auxiliary gene might be very occasional, such as the appearance of an antibiotic that might not be encountered again for a long time. Can bacteria prevent complete plasmid loss during long periods between selection events?

Several mechanisms are known to promote long-term plasmid persistence, including fitness-enhancing interactions between plasmids [8, 9], fitness-enhancing compensatory evolution of host chromosome and plasmid genes [10, 11, 12, 13], sufficiently frequent positive selection events [14], “weaponization” of the plasmid against competitors [15], and sustained beneficial plasmid effects for a fraction of the host population [16]. All of these mechanisms succeed by decreasing the cost of the plasmid or by providing sufficiently sustained positive selection. Here, we consider a worst-case scenario: a plasmid that provides only infrequent benefits and imposes high costs that cannot be alleviated by mutations without also losing access to the benefits (i.e. intrinsically costly). Is long-term persistence possible for such a plasmid?

To survive, plasmids have evolved mechanisms that prevent loss from hosts, especially during cell division. For example, plasmids can tune their copy number in the cell [17], actively partition plasmids between daughter cells [18], and deploy addiction systems to kill cells that do not retain the plasmid [19]. Thus, the probability of plasmid loss after cell division is reported as low as 10^*−*7^ [20]. Plasmids also spread horizontally from donor bacteria to acceptor bacteria via conjugation [21]. This exchange of genetic material is facilitated by some Type IV secretion systems (T4SS) [22]. However, the ubiquity of these extracellular structures makes them attractive receptors for phages, specifically plasmid-dependent or “male specific” phages that target conjugation machinery, especially the conjugative pilus [23]. Early studies suggested that conjugation rates were too slow to sustain costly plasmids even against small loss rates [24, 25, 26], but evidence is accumulating for sufficiently high conjugation rates, even in lab cultures [27, 28, 29, 30]. Are the mechanisms for plasmid spread and persistence sufficient to enable the long-term survival of plasmids that carry high-cost genes?

Previous studies have shown that high-conjugation rates are sufficient for the survival of high-cost plasmids, assuming a large supply of recipient cells to become transconjugants [31, 25, 32, 33], but this is not the only possible scenario. Thus, to complete our worst-case scenario, we demand that the plasmid must survive in the presence of competitors that are immune to (cannot receive) the plasmid, and may therefore outcompete plasmid-susceptible cells over time. Such plasmid-immune competitors could include bacteria with entry exclusion mechanisms [34], restriction enzymes targeting the costly plasmid [35, 36], or even archaeal or eukaryotic competitors. We expect such competing populations to be common in nature.

Population modeling of plasmids and their hosts has been a useful tool for understanding cell growth and plasmid abundance in experiments [37, 38], establish criteria for plasmid fixation in a population [31, 25, 32], and more [39, 40]. Here, we use this modeling framework to study the fate of high-cost plasmids in the presence of plasmid-dependent phages. We find that (1) horizontal transfer and persistence mechanisms are alone unlikely to enable the long-term survival of intrinsically high-cost plasmids in the presence of competitors that cannot receive the plasmid. In these conditions, we further find that (2) plasmid-dependent phages can – counterintuitively – promote the long-term survival of high-cost plasmids. Our results are robust within realistic parameter ranges and yield testable predictions.

Our models combine routine approaches to plasmid conjugation kinetics [31, 31, 32, 39] with Lotka-Volterra style predation dynamics describing the plasmid-dependent phage [33] and include delays between infection and lysis and between plasmid acquisition and pilus expression. Our models neglect demographic fluctuations, as they are negligible for systems with even moderate volumes (∼ 1 ml, see the Supplementary Materials for model details).

### Long-term survival of intrinsically costly plasmids is unexpected

High-cost plasmid genes face seemingly insurmountable odds against long-term survival. We first discuss this survival qualitatively and then quantitatively evaluate the timescale for high-cost plasmid loss within a population model. Finally, we demonstrate how plasmid-dependent phages can rescue high-cost plasmids from rapid extinction.

High-cost genes carried on plasmids are in constant danger of extinction. If horizontal transmission of a costly plasmid is too slow to overcome loss rates (Figure 1A, left), plasmid extinction is guaranteed unless the plasmid harbors other genes that provide a benefit greater than the cost. In this symbiotic regime, selection will drive the loss of costly genes, while beneficial genes are expected to migrate to the host chromosome [26]. Rapid horizontal transfer is also insufficient to save the costly gene(s). In the high-conjugation regime (Figure 1A, right), the plasmid will quickly spread to fixation as a parasite within the susceptible population. However, this rapid spread ultimately undermines plasmid survival because the plasmid cost leaves the host uncompetitive against plasmid-immune populations.

**Figure 1.**
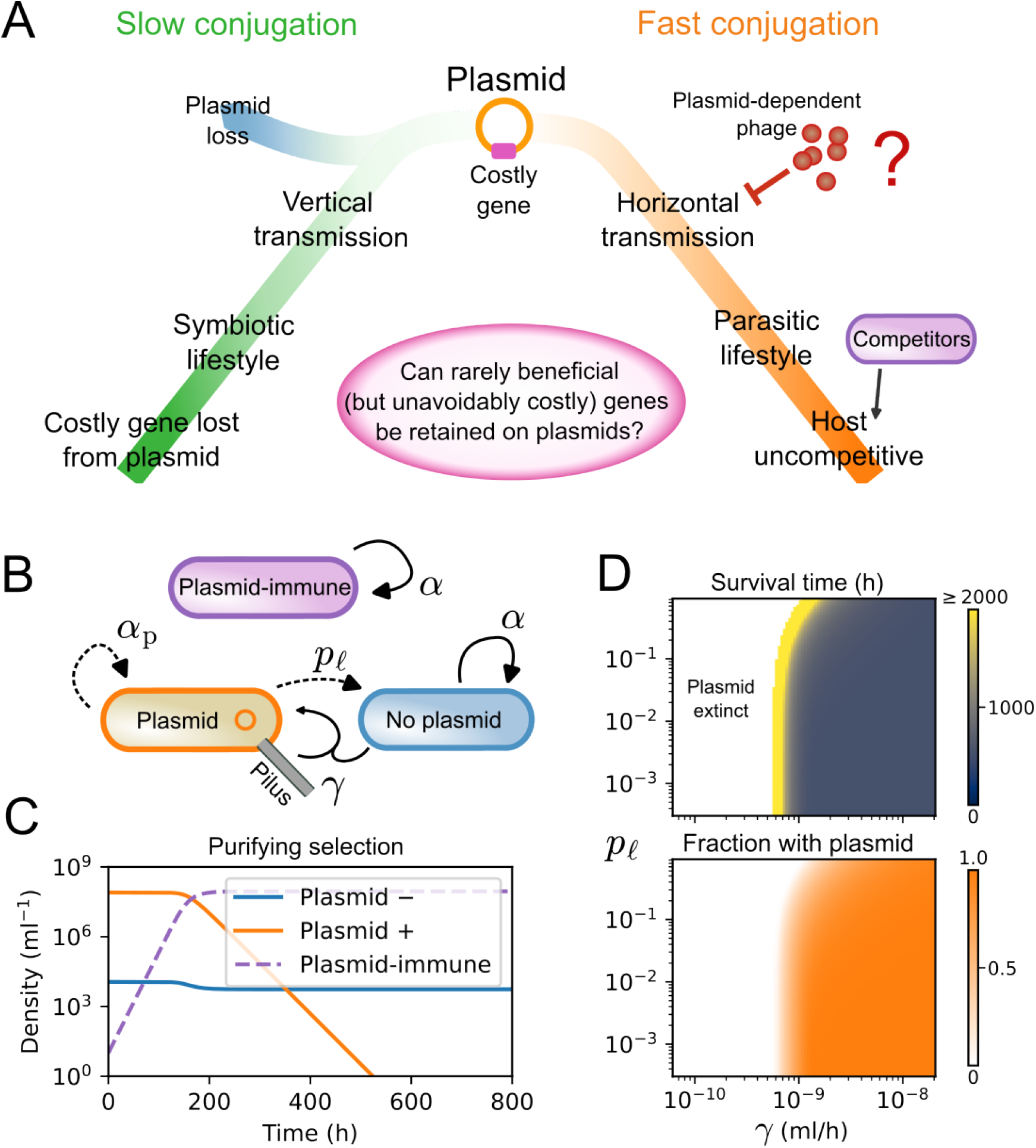
Can bacterial communities retain costly genes on plasmids until the genes become useful? (A) When horizontal transfer via conjugation is slow (left side in green), plasmid survival depends on vertical transmission (symbiosis) and costly genes are purified out of the plasmid. If conjugation is frequent instead, plasmid takeover can render the host uncompetitive (right side in orange). However, plasmid-dependent phages may disrupt this pathway. (B) A model ecology with a single costly plasmid that spreads quickly via conjugation (rate *γ*_hi_ = 10^*−*8^ ml hr^*−*1^) but cannot infect a subpopulation of cells (plasmid-immune) and is lost upon cell division with probability *p*_*l*_ = 10^*−*3^. Plasmid-containing cells grow at a rate *α*_*p*_ = 0.5/h that is 50% slower than plasmid-free cells *α* = 1.0/h. All growth is logistic with a carrying capacity *Q* = 10^8^/ml and a death/washout rate of *δ* = 0.1/hr. (C) At equilibrium (*t* = 0) plasmid-containing cells (orange) dominate the susceptible population (blue = plasmid free). The plasmid-immune competitor (dashed purple) is introduced and drives the plasmid to extinction, which occurs when abundance drops below 1.0/ml. (D) The survival time (top) of the plasmid following invasion by the competitor is always small, unless the conjugation and plasmid-loss rates are extremely finely balanced to produce an initial steady state (bottom) with only a small fraction of plasmid containing cells.

To estimate the extinction timescales for costly plasmids, we employ a population model (illustrated in Figure 1B) of a plasmid-susceptible population under invasion by a plasmid-immune competitor. Plasmid-free and plasmid-immune cells both grow logistically at a maximum rate of *α* = 1.0/hr, describing an ecology that is competitive in the absence of the plasmid. The maximum growth rate for plasmid-containing cells is *α*_*p*_ = (1 − Δ)*α* where Δ = 0.5 is the plasmid growth cost. This 50% growth penalty describes a very costly plasmid. The plasmid can spread horizontally (conjugation rate *γ*) and is lost upon cell division with probability *p*_*l*_ = 10^*−*3^. See Methods and SI Appendix for details.

When the conjugation rate is high enough, the plasmid can initially fix in the susceptible population. Starting with an invasion by a plasmid-immune competitor (*t* = 0, Figure 1C), within ∼ 500 h, the plasmid-immune population drives the plasmid to extinction (extinction threshold 1*/*ml), allowing the susceptible population to continue but without any long-term benefits the plasmid may carry. In models without plasmid-immune competitors [41, 32] and (presumably) experiments on lab cultures [29], long-term persistence can be sustained by high conjugation rates alone. However, we expect plasmid-immune competitors to be ubiquitous in natural environments.

Figure 1D plots the time to extinction as a function of *γ* and *p*_*l*_. The survival time of the plasmid is very short, except for a narrow region where the conjugation and loss probabilities are extremely finely balanced such that plasmid-containing cells constitute only a small fraction of the susceptible population at steady state (*t* = 0). In Figure 1D, the timescale for extinction increases as the the plasmid-containing fraction decreases because the impact of the plasmid scales with the plasmid-containing fraction. More precisely, the plasmid-susceptible cells (both plasmid-containing and plasmid-free) grow subject to the effective growth cost (SI Appendix 1.2)

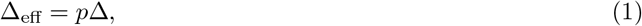

where *p* is the plasmid-containing fraction and Δ is the growth penalty of the plasmid. In the next section, we show that a plasmid-dependent phage can promote the survival of costly plasmids by keeping the plasmid-containing fraction low (and therefore reducing the impact of the plasmid on the population).

### Predation-enabled plasmid persistence

Plasmid-dependent phages target conjugative machinery (especially conjugative pili) as surface receptors, and therefore may achieve a broad host range [42], up to the range of plasmids they target. We focus specifically on lytic plasmid-dependent phages, many of which have recently been discovered [43, 44]. By introducing a plasmid-dependent phage into our population model (illustrated in Figure 2A) we demonstrate how predation by these phages can dramatically extend survival of high-cost plasmids. We refer to this dynamic as “predation-enabled plasmid persistence”.

**Figure 2.**
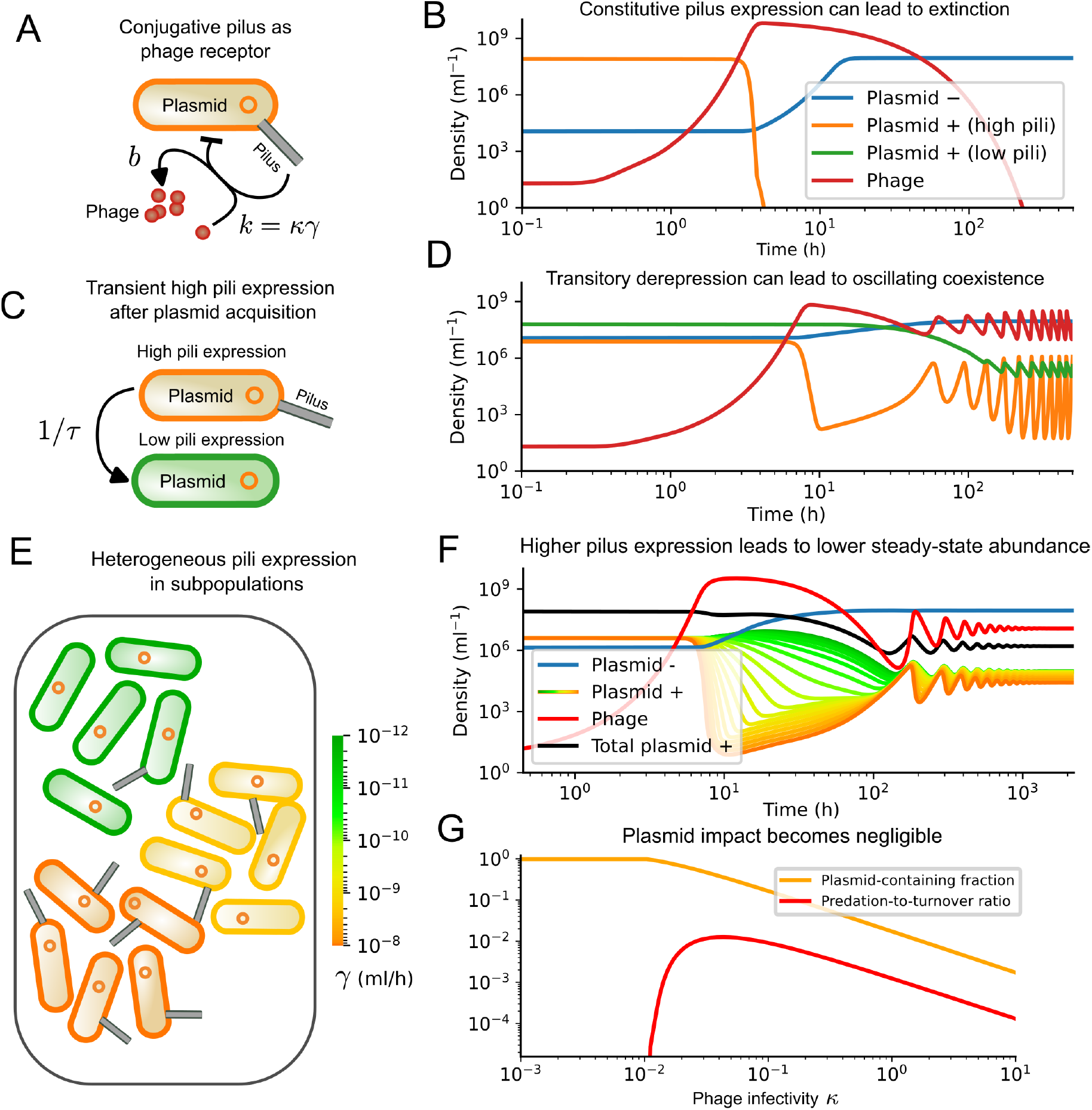
When plasmid-dependent phages are present, costly plasmids are maintained at low abundance. (A) Plasmid-dependent phages target conjugative pili as surface receptors. We assume that the phage absorption rate *k* and the conjugation rate *γ* are proportional (*k* = *κγ*). (B) When conjugative pili are constitutively expressed (conjugation rate *γ*_hi_ = 10^*−*8^ ml/hr, *κ* = 1), phage invasion can result in plasmid extinction (occurring when populations drop below 1/ml). To avoid extinction, a simple pilus regulatory strategy is to (C) only transiently express pili at a maximum level (*γ*_hi_) for an average time *τ* after plasmid acquisition, then dramatically reduce expression (*γ*_lo_ = 10^*−*12^ ml/hr). This regulation (D) prevents plasmid extinction but can lead to oscillations. Heterogeneity often dampens such oscillatory dynamics in nature, and we describe such heterogeneity by dividing the population into *N* = 20 subpopulations with constitutive expression of conjugation machinery to cover the range *γ*_lo_ ≤ *γ* ≤ *γ*_hi_. (F) Heterogeneity dampens the oscillations, with high-expression cell populations driven to lower abundance than low-expression populations. (G) As the phage infectivity (*κ*) increases above the phage survival threshold (*κ*≈10^*−*2^), the plasmid-containing fraction (orange) falls by orders-of-magnitude. Counterintuitively, losses due to phage predation also decrease (red, showing the ratio of phage infection to death/washout rate without the phage).

To describe the phage population in our model, we use dynamics that are routinely used in other contexts to model phage populations (see SI Appendix). Since the expression of conjugation machinery determines both the conjugation and phage infection kinetics, we assume that the phage infection rate constant *k* and the conjugation rate constant *γ* are proportional [33] (*k* = *κγ*, where *κ* is the proportionality constant), using *k* and *γ* parameter values that are typical for conjugation and lytic phage infection kinetics (see Supplementary Materials). The proportionality holds when (1) the phage adsorption rate is proportional to the frequency of encounters between phage particles and cells expressing at least one pilus, and (2) the rate of transconjugant formation is proportional to the frequency of encounters between a plasmid-susceptible cell without the plasmid and a plasmid-containing cell expressing at least one pilus.

We first examine the dynamics of plasmid-dependent phages into a bacterial population with a single costly plasmid, without plasmid-immune competitors (Figure 2A). In the invasion of Figure 2B, expression of conjugation machinery is constitutive such that conjugation is always occurring at the maximum rate (see Methods). The model and conditions are the same as in Figure 1C, but with the addition of the phage. In this case, the plasmid is driven to extinction by the phage. However, such extinction can be avoided when expression of the conjugation machinery is regulated [21] (not constitutive) or heterogeneous within the population [33].

First, we consider phage invasion when conjugation is regulated through transitory de-repression [27], illustrated in Figure 2C. In this mechanism, transconjugants express conjugation machinery at high levels (achieving a fast conjugation rate *γ*_hi_) for only an average time *τ* before repressing the machinery permanently to low levels (conjugation rate *γ*_lo_). Thus, conjugation is primarily driven by recent transconjugants. Even this simple regulatory framework prevents extinction is prevented and plasmid abundance is maintained at a low level, even as *γ*_lo_ and *τ* vary over orders-of-magnitude (see Figure S1). In the limit-cycle dynamics of Figure 2D, the population of plasmid-containing cells oscillates with frequency 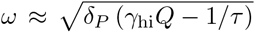 around a fixed point with a derepressed population ≈ *δ*_P_*/*(*bκγ*_hi_), where *δ*_P_ is the death (washout) rate of the phage and *b* is the lysis burst size (see SI Appendix 2.1).

Next, we consider a population with heterogeneous expression of conjugation machinery, as illustrated in Figure 2E. Such heterogeneity can result from varying cell activity [45], plasmid “interference” [46], quorum sensing [47], and more [21]. To approximate these behaviors, we evenly bin (on a log scale) the initial plasmid-susceptible population by average conjugation rate *γ* into *N* = 20 groups that vary from low to high pili expression (conjugation rate). Within each group, cells either have the plasmid or not. As a result of this heterogeneity, the oscillations in plasmid abundance are dampened and a steady state is reached after a few hundred hours with High-conjugation cells suppressed to lower abundance. Previous modeling work demonstrated that heterogeneity can prevent plasmid extinction by phages [33], but did not consider how phages themselves might promote plasmid survival.

The impact of plasmid-dependent phages on costly plasmids in the above examples suggests a generalization with potentially far-reaching consequences: after initial transient dynamics, plasmid-dependent phages pin down the abundance of plasmid-containing cells to low levels, but not to extinction. This drives down the effective growth cost Δ_eff_ of the plasmid to the susceptible population (Equation 1). However, this raises another question – to what extent is the effective fitness boost to the bacteria due to smaller growth costs counteracted by an effective fitness decrease driven by cell losses via phage predation?

To evaluate the impact of predation losses on the plasmid-susceptible population, we evaluate the steady-state behavior of Figure 2F as a function of the phage infectivity *κ*. Figure 2G shows both the plasmid-containing fraction (orange) and the ratio *ρ* (red)

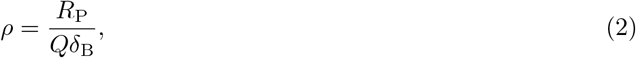

where *R*_P_ is the total rate of phage infections across the susceptible population, and *Qδ*_B_ is the death (washout) rate of the bacterial population when it fills the carrying capacity *Q*. Therefore, *ρ* captures the relative magnitudes of cell loss due to death (washout) versus phage predation. At low infectivity, the phage population cannot be sustained and goes extinct, allowing the plasmid to spread to fixation in the population. As the infectivity *κ* increases to a value typical for lytic phages (*κ* ∼ 1 or higher), the plasmid-containing fraction falls dramatically to well below a percent. Counterintuitively, the rate of phage predation *ρ* also decreases by orders of magnitude because predation limits prey abundance. For typical infection rates of lytic phages (*k*_hi_ = 10^*−*7^ ml/h or *κ* = 10), the phage negligibly increases the total turnover rate of the plasmid-susceptible population.

The results in Figure 2 demonstrate that plasmid-dependent phages can in principle confine high-cost plasmids to low abundance – but not extinction – without any fine tuning or special regulation, and simultaneously inflict minimal predation losses as a result. This regime is precisely where persistence of the costly plasmid is maximized. Figure 3 illustrates how phage predation dramatically increases costly plasmid survival times.

**Figure 3.**
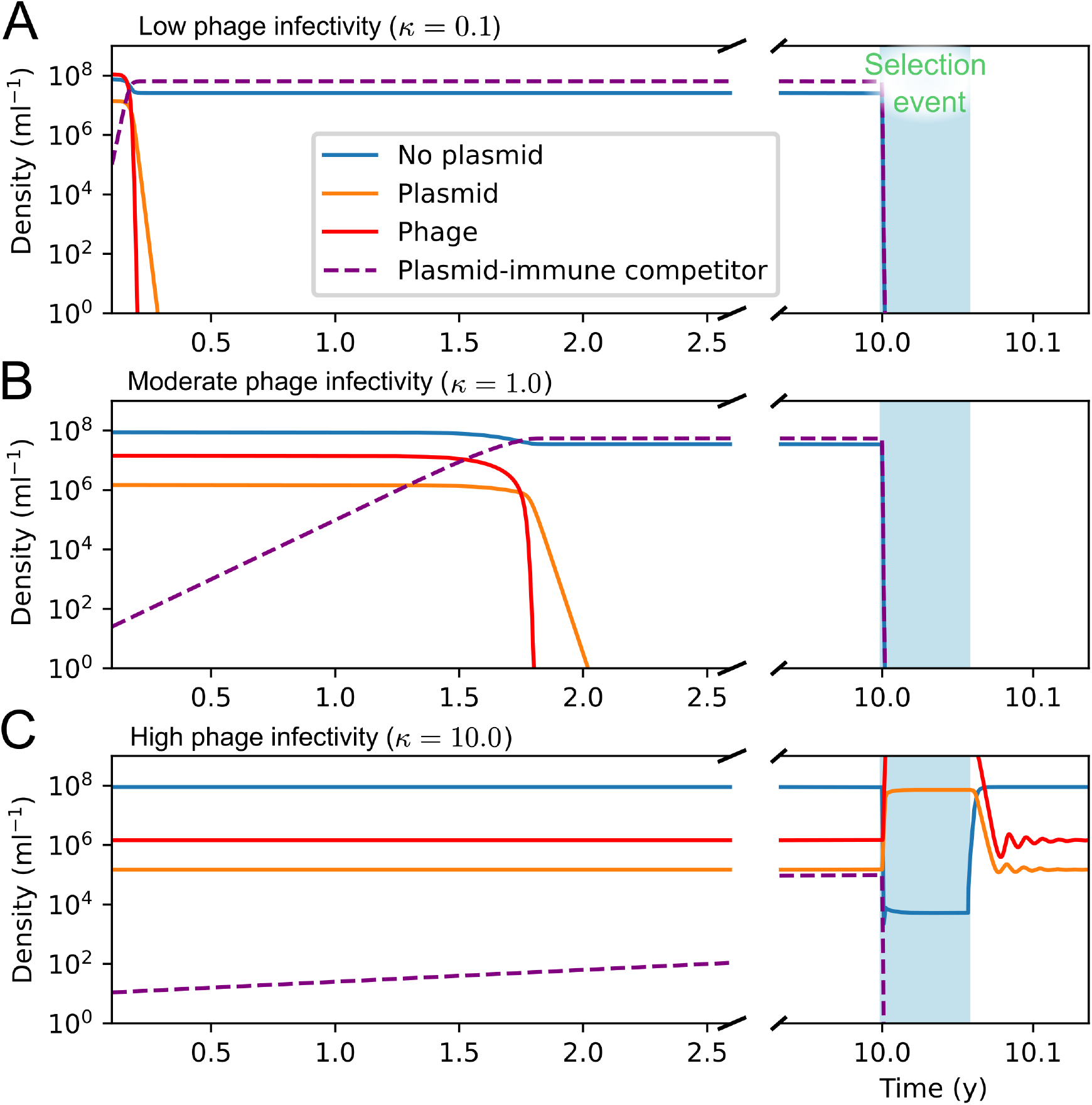
Plasmid-dependent phages can enable costly plasmids to survive between rare selection events. In this example, survival of the bacteria requires the persistence of a costly antibiotic resistance plasmid until the appearance of an antibiotic at *t* = 10 years (selection event, shown in light blue). The antibiotic kills plasmid-free cells at a rate 2.0/h. (A) Even at low infectivity *κ* = 0.1 (*k*_hi_ = *κγ*_hi_ = 10^*−*9^ ml/h) the plasmid persists for ∼ 2000 h, much longer than in the absence of the phage (Figure 1C), but far too short to reach the selection event, so all cells as well as the plasmid and phage die. As phage infectivity increases from (B) *κ* = 1.0 (*k*_max_ = 10^*−*8^ ml/h) to (C) *κ* = 10.0 (*k*_max_ = 10^*−*7^ ml/h), the costly plasmid imposes progressively less and less of a burden on the host population, slowing the timescale of plasmid extinction by years. In (C), the plasmid survives long enough to provide protection from the antibiotic. The model and other parameter values are the same as in Figures 1C, 2F, and 2G.

In the example of Figure 3, the plasmid-susceptible population must hold onto a high-cost plasmid for 10 years, after which a plasmid selection event occurs (i.e. the appearance of an antibiotic). During this selection event, cells without the plasmid are killed at a rate *r*_s_ = 2.0/h. If the plasmid survives until the event, it can rescue the susceptible population and ultimately allow the susceptible population to outcompete the plasmid-immune competitor that cannot receive protection from the plasmid. As phage infectivity is increased from low (*κ* = 0.1) to high (*κ* = 10.0, still within typical ranges, see Methods), the timescale for invasion of the plasmid immune competitor and subsequent plasmid extinction is increased from ∼ 1000 h to many years (*κ* = 10.0). When the selection event occurs, the costly plasmid quickly switches to a vertical-transmission lifestyle.

The simulation of Figure 3 is a simplification of ecological dynamics in nature and is only intended to demonstrate that long-term persistence of unavoidably high-cost plasmids can surprisingly be enabled by phages, even over timescales of years. Over such a period, ecological and evolutionary forces may drive the formation of new hosts, plasmid descendants, and even competitors. However, in the absence of the phage, the high-cost plasmid only survives for a short period (Figure 1C), making its long-term evolution impossible. Plasmid survival rates after the selection event also may vary because the high-conjugation populations are not needed during the selection event may therefore be lost. Replacements might arise through mutations in the regulation of conjugative machinery or changes in cell state, but may not always emerge again. In Figure 2E-G, we consider slow interconversion of conjugation states (*T*_mix_ = 100 h), reflecting drift in the expression of conjugation machinery over time (otherwise critical high-conjugation populations are lost during the selection event, see Figure S2). None of the above considerations impact our main conclusion: plasmid-dependent phages can enable the survival of intrinsically costly plasmids for long periods between selection events.

## Discussion

We propose that plasmid-dependent phages can, counterintuitively, promote the survival of their target plasmids by suppressing plasmids to low, but nonzero, abundance within their host populations. We observed this behavior in population models with typical parameter values (and over a broad range, see Figures 2-3 and Figure S1). Predation-enabled plasmid persistence can occur alongside other mechanisms shown to promote plasmid survival, but distinguishes itself by enabling the survival of plasmids with unavoidable high costs and in the presence of competitors that cannot receive the plasmid, a remarkable feat.

Predation-enabled persistence of high-cost plasmids in our models relies on two key features: (1) the presence of plasmid-immune competitors, and (2) the negative feedback inherent in predator-prey dynamics. With plasmid-immune competitors, intrinsically costly plasmids can only survive for long periods if the conjugation and loss kinetics are unrealistically finely balanced to maintain the plasmid at low abundance (Figure 1D). While previous analyses considered plasmid-dependent phages to only decrease the likelihood of plasmid persistence [33, 48], we reach a different conclusion. By predating on cells expressing conjugative pili, plasmid-dependent phages can establish a critical negative feedback: as plasmid abundance increases, so does phage predation. When plasmid abundance falls, phage predation also falls, resulting in suppressed high-cost plasmid populations, precisely the conditions required for costly-plasmid survival (Figure 3). Non-lytic plasmid-dependent phages may also constrain the spread of high-cost plasmids, such as filamentous phages that spread by secretion of phage particles from infected hosts instead of lysing their hosts[49]. Diverting cellular resources to produce phage particles (or occlusion of the conjugative pilus by phage [50]) may limit the conjugation rates of host bacteria, leading to falling abundance of plasmid-containing cells.

Our theory yields at least three testable predictions: (1) Intrinsically high-cost plasmids may exist in nature where plasmid-dependent phages are found. These plasmids must be sustained by tiny bacterial subpopulations with high conjugation rates. Previous analyses considered such “super donor” populations [28, 33, 41], but did not consider whether predation-enabled plasmid persistence may self-regulate these populations to low levels. (2) Introducing plasmid-dependent phages to cultures where they are absent drive an increase in average bacterial fitness (such as growth rates), by cutting down the abundance of any costly plasmids. (3) Plasmid-dependent phages can be used as a capture reagent for rare and costly plasmids, which we discuss next.

Our findings suggest that high-risk, high-reward genes can be maintained at low abundance where low-depth metagenomics approaches may fail to capture them. Because high-cost plasmids are forced to express conjugation machinery at high levels to survive, plasmid-dependent phages themselves might be used for high-cost plasmid discovery. For example, lysis-deficient plasmid-dependent phages could be used to immobilize otherwise flowing cells that express conjugation machinery. This would create a sample of bacteria greatly enriched for plasmids in this low-abundance reservoir. Already, similar experiments have demonstrated trapping of bacteria expressing T4 receptors [51, 52]. Alternatively, plasmid-dependent phage vectors [53] could be used to deliver antibiotic resistance markers to cells expressing conjugation machinery, allowing these plasmid-containing cells to be selected out of a population. Significant revisions may be forthcoming to our understanding of plasmid-dependent phages and their role in natural bacterial communities.

## Data Availability

The code used to perform all analyses is archived through Zenodo[54], and is available at GitHub at https://github.com/JYuly/Plasdepphage-eco.

## Acknowledgements

This work was supported in part by the Peter B. Lewis Lewis-Sigler Institute/Genomics Fund through the Lewis-Sigler Institute for Integrative Genomics at Princeton University. This work was also supported in part by Princeton University through the Center for the Physics of Biological Function. This work was also supported by the National Institutes of Health under Award No. R35GM156427. The content is solely the responsibility of the authors and does not necessarily represent the official views of the National Institutes of Health. We thank Qiwei Yu and Colin Scheibner for helpful discussions.

