## Supplemental Information for "Do plasmid dependent phages promote the survival of costly plasmids?"

### Supporting Information for: Can plasmid-dependent phages promote the survival of costly plasmids?

**3 Model with heterogeneous expression of conjugation machinery** **5**

### 1 Population model with single mode of pilus expression

We consider the ecological dynamics of a community of plasmid-dependent phages (abundance $P$), bacteria containing the plasmid ($B_\mathrm{p}$), without the plasmid ($B_0$), and a competitor that is immune to the plasmid ($B_\mathrm{c}$). To describe this ecology, we combine routine approaches to plasmid conjugation dynamics [1, 2, 3] and predation dynamics to describe the plasmid-dependent phage [4]. When all plasmid-containing cells express the pilus at the same rate, we describe the dynamics using the delay integro-differential equations that describe bacterial growth, conjugation, and phage infection with delays $T_\mathrm{pili} = T_\mathrm{lysis} = 0.5$ h between receiving the plasmid and expressing conjugation machinery and from phage infection to lysis, respectively. The equations are

$$\frac{\mathrm{d}B_0(t)}{\mathrm{d}t} = \left[R_0(t) - \delta_\mathrm{B}\right] B_0(t) + p_l R_\mathrm{p}(t) B_\mathrm{p}(t) - \gamma B_\mathrm{p}(t) B_0(t), \tag{1}$$

$$\frac{\mathrm{d}B_\mathrm{p}(t)}{\mathrm{d}t} = \left[(1-p_\ell) R_p(t) - \delta_\mathrm{B}\right] B_\mathrm{p}(t) - k B_\mathrm{p}(t) P(t) + \gamma e^{-T_\mathrm{pili}\delta_\mathrm{B}}\; B_\mathrm{p}(t - T_\mathrm{pili})\; B_0(t - T_\mathrm{pili}), \tag{2}$$

$$\frac{\mathrm{d}P(t)}{\mathrm{d}t} = b k e^{-T_\mathrm{lysis}\delta_B}\; B_\mathrm{p}(t - T_\mathrm{lysis}) P(t - T_\mathrm{lysis}) - \delta_\mathrm{P} P(t) - k P(t) \int_0^{T_\mathrm{lysis}} \mathrm{d}u\;\, k P(t-u) B_\mathrm{p}(t-u) e^{-\delta_\mathrm{B} u}, \tag{3}$$

$$\frac{\mathrm{d}B_\mathrm{c}}{\mathrm{d}t} = R_\mathrm{c} B_\mathrm{c}(t) - \delta_\mathrm{B} B_\mathrm{c}(t), \tag{4}$$

where $\delta_\mathrm{B}$ and $\delta_\mathrm{P}$ are the bacterial and phage death (washout) rates, $p_\ell$ is the probability of plasmid loss upon cell division. The conjugation rate is $\gamma$ and the phage infection (absorption) rate is $k = \kappa\gamma$. The $R_i$ are the instantaneous growth rates for the bacterial populations ($i = 0, \mathrm{p}, \mathrm{c}$ or plasmid-free, plasmid containing, and competitor, respectively). We model the growth of these populations as logistic, with a growth cost $\Delta$ for bacteria carrying the plasmid

$$R_0(t) = R_\mathrm{c}(t) = \alpha_0 \left(1 - \frac{B_0(t) + B_\mathrm{p}(t) + B_\mathrm{c}(t) + B_\mathrm{I}(t) + B_\mathrm{nt}(t)}{Q}\right) = \alpha_0 \left(1 - \frac{B_\mathrm{T}}{Q}\right), \tag{5}$$

$$R_{\mathrm{p}}(t) = \alpha_{\mathrm{p}} \left(1 - \frac{B_{\mathrm{T}}}{Q}\right), \tag{6}$$

where $\alpha_{\mathrm{p}} = (1-\Delta)\alpha_0$, and $B_{\mathrm{T}}$ is the sum of all of the bacterial populations. The parameters $k$, $b$, $\delta_{\mathrm{P}}$, and $\delta_{\mathrm{B}}$ are the phage absorption (infection) rate, the phage burst size, and the phage and bacterial death (washout) rates. Also, we use $k = \kappa\gamma$ and $\delta_{\mathrm{P}} = \delta_{\mathrm{B}}$ as mentioned in the main text. The $B_{\mathrm{I}}$ and $B_{\mathrm{nt}}$ populations are infected and recent transconjugants, respectively, that have not yet burst or expressed conjugation machinery:

$$B_{\mathrm{I}} = \int_0^{T_{\mathrm{lysis}}} \mathrm{d}u \;\; kP(t-u)B_{\mathrm{p}}(t-u)e^{-\delta_{\mathrm{B}}u}, \tag{7}$$

$$B_{\mathrm{nt}} = \int_0^{T_{\mathrm{pili}}} \mathrm{d}u \;\; \gamma B_0(t-u)B_{\mathrm{p}}(t-u)e^{-\delta_{\mathrm{B}}u}, \tag{8}$$

A similar approach (but without delays) was previously used to model plasmid-dependent phages [4].

Unless otherwise stated, we use $\alpha = 1.0$/h, $\alpha_p = 0.5$/h, $Q = 10^8$/ml, $p_\ell = 10^{-3}$, and $\kappa = 1$. In all simulations, we set the extinction threshold to 1.0/ml. To estimate the value of the phage infectivity parameter $\kappa$, we assume that the phage infection kinetics correspond to those typical for lytic phages. Specifically, pilus expression for the F plasmid can be permanently derepressed to yield a maximum conjugation rate of around $\gamma_{\mathrm{hi}} \approx 2\times10^{-9}$ ml/hr [5] to $3\times10^{-8}$ ml/h [6]. Thus, we use $\gamma_{\mathrm{hi}} \sim 10^{-8}$ ml/hr, although conjugation rates as high as $10^{-6}$ ml/hr have been reported for some plasmids [7]. We expect that phages absorb to these fully derepressed cells at rates typical for lytic phages. Phage absorption rates are typically $k \sim 10^{-7}$ ml/h or less [8]. Thus, the "infectivity constant" $\kappa$ in our model should be $\kappa = k/\gamma \sim 10^1$ or less. Unless stated otherwise, we conservatively use $\kappa = 1$. A value $\kappa \approx 10^{-4}$ was previously suggested [4], but such infection rates would be anomalously low among lytic phages, was derived assuming homogeneity in experimental bacterial populations, and predicts phage extinction where it was not observed [9]. During the selection events of Figure 3, all cells without the plasmid die at a rate 2.0/h.'

Suppression of costly plasmids to low abundance in our model raises the possibility that demographic stochasticity might become important. However, absolute frequency – not population density – sets the scale for demographic fluctuations. For example, if the system in Figure 3 has a volume $V \sim 1$ mL, it would contain $\sim 10^5$ plasmid-containing cells at steady state, so demographic stochasticity would be negligible (in tiny microenvironments with volume $V \sim 0.01$ mL or less, demographics fluctuations may be important).

All numerical calculations were performed using the automatic stiffness detection method [10] implemented in the SciPy [11] wrapper of ODEPACK [12].

#### 1.1 Approximating delays with intermediate states

Delay equations like those above can be unwieldy. Fortunately, we approximate the delays for pilus expression and lysis using a series of $M$ intermediate cell states

$$\text{Conjugation event } \to S_1 \to S_2 \to \ldots \to S_M \to \text{Transconjugant expresses conjugation machinery}, \tag{9}$$

$$\text{Infection event } \to I_1 \to I_2 \to \ldots \to I_M \to \text{Lysis}. \tag{10}$$

This approximation has been used previously to describe lytic phages [13]. Thus, we approximate the delay equations in the previous section using the ordinary ODEs

$$\frac{\mathrm{d}B_0}{\mathrm{d}t} = (R_0 - \delta_{\mathrm{B}})\,B_0 + p_\ell R_{\mathrm{p}} B_{\mathrm{p}} - \gamma B_{\mathrm{p}} B_0, \tag{11}$$

$$\frac{\mathrm{d}S_1}{\mathrm{d}t} = \gamma B_0 B_{\mathrm{p}} - \left(\frac{M}{T_{\mathrm{pili}}} + \delta_{\mathrm{B}}\right) S_1, \tag{12}$$

$$\frac{\mathrm{d}S_i}{\mathrm{d}t} = \frac{M}{T_{\mathrm{pili}}} S_{i-1} - \left(\frac{M}{T_{\mathrm{pili}}} + \delta_{\mathrm{B}}\right) S_i \quad (1 < i \le M), \tag{13}$$

$$\frac{\mathrm{d}B_{\mathrm{p}}}{\mathrm{d}t} = \left[R_{\mathrm{p}}(1-p_\ell) - \delta_{\mathrm{B}}\right] B_{\mathrm{p}} + \frac{M}{T_{\mathrm{pili}}} S_M - kB_{\mathrm{p}}P, \tag{14}$$

$$\frac{\mathrm{d}I_1}{\mathrm{d}t} = kB_{\mathrm{p}}P - \left(\frac{M}{T_{\mathrm{lysis}}} + \delta_{\mathrm{B}}\right) I_1, \tag{15}$$

$$\frac{\mathrm{d}I_i}{\mathrm{d}t} = \frac{M}{T_{\mathrm{lysis}}} I_{i-1} - \left(\frac{M}{T_{\mathrm{lysis}}} + \delta_{\mathrm{B}}\right) I_i \quad (1 < i \le M), \tag{16}$$

$$\frac{\mathrm{d}P}{\mathrm{d}t} = b\frac{M}{T_{\mathrm{lysis}}}I_M - \delta_{\mathrm{P}}P - kP\sum_{i=1}^{M}I_i, \tag{17}$$

where we have dropped the explicit time dependence because there are no longer delays in the equations. In all simulations, we choose $M = 5$. As before, the intermediate populations do not divide but do contribute to the logistic term

$$R_{0,p} = \alpha_{0,p}\frac{1}{Q}\left(Q - B_0 - B_{\mathrm{p}} - \sum_{i=1}^{M}(S_i + I_i)\right). \tag{18}$$

#### 1.2 Effective plasmid growth cost in susceptible populations

To calculate the growth rate $R_{\mathrm{s}}(t)$ of the plasmid-susceptible population $B_{\mathrm{s}} = B_0 + B_{\mathrm{p}}$, we first add the growth terms of the plasmid-free $B_0$ and plasmid-containing cells

$$\begin{aligned} R_{\mathrm{s}}B_{\mathrm{s}} &= (R_0 - \delta_B)B_0 + (R_{\mathrm{p}} - \delta_{\mathrm{B}})B_{\mathrm{p}} \\ &= \left[(R_0 - \delta_{\mathrm{B}})(1-p) + ((1-\Delta)R_0 - \delta_{\mathrm{B}})p\right]B_{\mathrm{s}} \\ &= [(1 - p\Delta)R_0 - \delta_{\mathrm{B}}]B_{\mathrm{s}}, \end{aligned} \tag{19}$$

where $p = B_{\mathrm{p}}/B_{\mathrm{s}}$ is the plasmid-containing fraction of plasmid-susceptible cells. Thus, we identify $\Delta_{\mathrm{eff}} = p\Delta$ as the effective growth cost of the plasmid on the plasmid-susceptible population.

### 2 Model with transitory derepression of conjugation

To capture transitory derepression dynamics in Figure 2D, we introduce two types of plasmid containing cells. One species $B_p^{\mathrm{hi}}$ expresses conjugation machinery at high levels to achieve a conjugation rate $\gamma_{\mathrm{hi}}$, and the other $B_p^{\mathrm{lo}}$ at low levels with conjugation rate $\gamma_{\mathrm{lo}}$:

$$\frac{\mathrm{d}B_0}{\mathrm{d}t} = (R_0 - \delta_{\mathrm{B}})B_0 + p_\ell R_{\mathrm{p}}(B_{\mathrm{p}}^{\mathrm{hi}} + B_{\mathrm{p}}^{\mathrm{lo}}) - \left(\gamma_{\mathrm{hi}}B_{\mathrm{p}}^{\mathrm{hi}} + \gamma_{\mathrm{lo}}B_{\mathrm{p}}^{\mathrm{lo}}\right)B_0, \tag{20}$$

$$\frac{\mathrm{d}S_1}{\mathrm{d}t} = \left(\gamma_{\mathrm{hi}}B_{\mathrm{p}}^{\mathrm{hi}} + \gamma_{\mathrm{lo}}B_{\mathrm{p}}^{\mathrm{lo}}\right)B_0 - \left(\frac{M}{T_{\mathrm{pili}}} + \delta_{\mathrm{B}}\right)S_1, \tag{21}$$

$$\frac{\mathrm{d}S_i}{\mathrm{d}t} = \frac{M}{T_{\mathrm{pili}}}S_{i-1} - \left(\frac{M}{T_{\mathrm{pili}}} + \delta_{\mathrm{B}}\right)S_i \qquad i \in \{1, 2, ..., M\}, \tag{22}$$

$$\frac{\mathrm{d}B_{\mathrm{p}}^{\mathrm{hi}}}{\mathrm{d}t} = \left[R_{\mathrm{p}}(1 - p_\ell) - \delta_{\mathrm{B}} - \frac{1}{\tau}\right]B_{\mathrm{p}}^{\mathrm{hi}} + \frac{M}{T_{\mathrm{pili}}}S_M - k_{\mathrm{hi}}B_{\mathrm{p}}^{\mathrm{hi}}P, \tag{23}$$

$$\frac{\mathrm{d}B_{\mathrm{p}}^{\mathrm{lo}}}{\mathrm{d}t} = \left[R_{\mathrm{p}}(1 - p_\ell) - \delta_{\mathrm{B}}\right]B_{\mathrm{p}}^{\mathrm{lo}} + \frac{1}{\tau}B_{\mathrm{p}}^{\mathrm{hi}} - k_{\mathrm{lo}}B_{\mathrm{p}}^{\mathrm{lo}}P, \tag{24}$$

$$\frac{\mathrm{d}I_1}{\mathrm{d}t} = \left(k_{\mathrm{hi}}B_{\mathrm{p}}^{\mathrm{hi}} + k_{\mathrm{lo}}B_{\mathrm{p}}^{\mathrm{lo}}\right)P - \left(\frac{M}{T_{\mathrm{lysis}}} + \delta_{\mathrm{B}}\right)I_1, \tag{25}$$

$$\frac{\mathrm{d}I_i}{\mathrm{d}t} = \frac{M}{T_{\mathrm{lysis}}}I_{i-1} - \left(\frac{M}{T_{\mathrm{lysis}}} + \delta_{\mathrm{B}}\right)I_i \qquad i \in \{1, 2, ..., M\}, \tag{26}$$

$$\frac{\mathrm{d}P}{\mathrm{d}t} = b\frac{M}{T_{\mathrm{lysis}}}I_M - \delta_{\mathrm{P}}P - kP\sum_{i=1}^{M}I_i, \tag{27}$$

$$R_0 = \alpha_0\frac{1}{Q}\left(Q - B_0 - B_{\mathrm{p}}^{\mathrm{hi}} - B_{\mathrm{p}}^{\mathrm{lo}} - \sum_{i=1}^{M}(S_i + I_i)\right). \tag{28}$$

$$R_{\mathrm{p}} = (1 - \Delta)R_0 \tag{29}$$

#### 2.1 Phage invasions with varying parameters in transitory derepression model

Figure S1 shows the results of the phage invasion in Figure 2D after 5000 h as the low conjugation rate $\gamma_{\mathrm{lo}}$ and repression time $\tau$ are varied by orders of magnitude. Green shows the plasmid-containing fraction $p$ and red the phage predation-to-turnover ratio $\rho = R_{\mathrm{P}}/Q\delta_B$ (Equation 2 in the main text) averaged over a time $t_{\mathrm{ave}} = 500$ h, thus over any oscillations that might persist after the invasion. Sometimes, the plasmid cannot survive on its own or phage invasion results in extinction of the plasmid (and therefore also extinction of the phage, shown in grey), but a wide range of values for these parameters results in coexistence.

As the infectivity $\kappa$ of the phage increases from small (left column) to large (right column), both $p$ and $\rho_{\mathrm{P}}$ become small regardless of the precise "strategy" (values of $\gamma_{\mathrm{lo}}$ and $\tau$) the plasmid uses for transitory derepression of its conjugation machinery. Thus, as long as the phage is sufficiently infectious ($\kappa \approx 1.0$ or greater), the plasmid can never take over the susceptible population. This reflects the unavoidable tradeoff between conjugation and predation: high-cost plasmids require high conjugation rates to survive, but this makes them vulnerable to the plasmid-dependent phage.

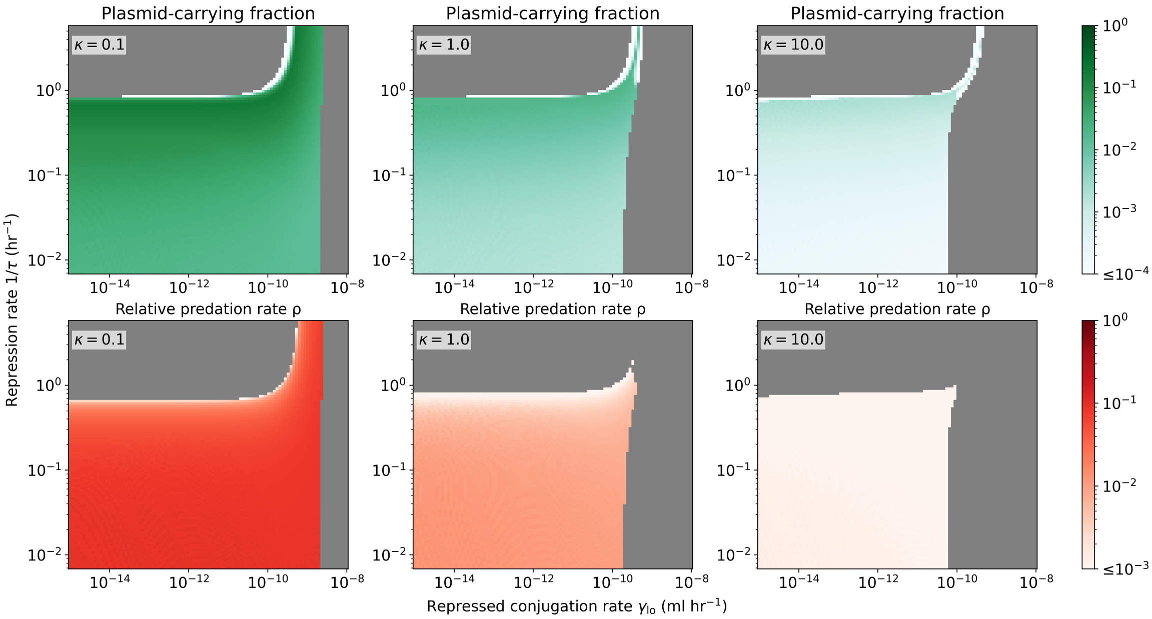

Figure S1: **Results of phage invasions for cells with transitory derepression of conjugation machinery.** Simulations of phage invasions like that in Figure 2D were performed for $t = 5000$ h, varying the parameters $\gamma_{\mathrm{lo}}$ and $\tau$ over several orders of magnitude. The top panels show the plasmid-containing fraction and the bottom panels the predation-to-turnover ratio $\rho$ (Equation 2 in the main text). As the phage infectivity increases from $\kappa = 0.1$ (left column) to $\kappa = 1.0$ (middle column) to $\kappa = 10.0$ (right column), both the plasmid-containing fraction and $\rho_{\mathrm{P}}$ become small.

#### 2.2 Fixed point analysis

In the oscillations of Figure 2D, the growth of the plasmid-containing cells is dominated by conjugation of recent transconjugants (derepressed cells) with conjugation rate $\gamma_{\mathrm{hi}}$ and the population of plasmid-free cells is approximately constant, just below the carrying capacity ($B_0(t) \approx Q$). Also, the phage growth is dominated by infections of derepressed cells because repressed cells (conjugation rate $\gamma_{\mathrm{lo}}$) rarely express conjugation machinery. Thus, for the oscillations in Figure 2D, the equations that describe the high-conjugation bacterial and phage population are

$$\frac{\mathrm{d}B^{\mathrm{hi}}(t)}{\mathrm{d}t} \approx \gamma_{\mathrm{hi}} e^{-T_{\mathrm{pili}}\delta_B}\ QB_{\mathrm{p}}^{\mathrm{hi}}(t - T_{\mathrm{pili}}) - \frac{1}{\tau}B_{\mathrm{p}}^{\mathrm{hi}}(t) - k_{\mathrm{hi}}B_{\mathrm{p}}^{\mathrm{hi}}(t)P(t), \tag{30}$$

$$\frac{\mathrm{d}P(t)}{\mathrm{d}t} \approx bk_{\mathrm{hi}}e^{-T_{\mathrm{lysis}}\delta_{\mathrm{B}}}\ B_{\mathrm{p}}^{\mathrm{hi}}(t - T_{\mathrm{lysis}})P(t - T_{\mathrm{lysis}}) - \delta_{\mathrm{P}}P(t) - k_{\mathrm{hi}}P(t)\int_0^{T_{\mathrm{lysis}}} \mathrm{d}u\ \ kP(t-u)B_{\mathrm{p}}^{\mathrm{hi}}(t-u)e^{-\delta_{\mathrm{B}}u}. \tag{31}$$

The nontrivial fixed point $B_p^{\mathrm{hi}*}, P^* \neq 0$ of this system is determined by the equations

$$\gamma_{\mathrm{hi}}e^{-T_{\mathrm{pili}}\delta_{\mathrm{B}}}\ Q - \frac{1}{\tau} - k_{\mathrm{hi}}P^* = 0, \tag{32}$$

$$bk_{\mathrm{hi}}e^{-T_{\mathrm{lysis}}\delta_{\mathrm{B}}}\ B_{\mathrm{p}}^{\mathrm{hi}*} - \delta_{\mathrm{P}} - \frac{k_{\mathrm{hi}}^2}{\delta_{\mathrm{B}}}P^*B_{\mathrm{p}}^{\mathrm{hi}*}\left(1 - e^{-\delta_{\mathrm{B}}T_{\mathrm{lysis}}}\right) = 0. \tag{33}$$

The above equations assumes that the transconjugants produced by the low-conjugation population $B_p^{\mathrm{lo}}$ are small, and that the low-conjugation population do not significantly absorb phage. This approximation is valid when $\gamma_{\mathrm{lo}}$ is small. The solution to the above equations is

$$B_{\mathrm{p}}^{\mathrm{hi}*} = \frac{\delta_{\mathrm{P}}}{bk_{\mathrm{hi}}e^{-T_{\mathrm{lysis}}\delta_{\mathrm{P}}} - \frac{k_{\mathrm{hi}}^2}{\delta_{\mathrm{B}}}P^*\left(1 - e^{-\delta_{\mathrm{B}}T_{\mathrm{lysis}}}\right)}, \qquad P^* = \frac{1}{k_{\mathrm{hi}}}\left(\gamma_{\mathrm{hi}}Qe^{-T_{\mathrm{pili}}\delta_{\mathrm{B}}} - \frac{1}{\tau}\right). \tag{34}$$

In our simulations, $\delta_P = \delta_B = 0.01/\mathrm{h}$ and $T_{\mathrm{lysis}} = T_{\mathrm{pili}} = 0.5$ h, so $e^{-\delta_B T_{\mathrm{lysis}}} = e^{-\delta_B T_{\mathrm{pili}}} \approx 0.995$. Therefore, we can approximate the fixed point as

$$B_{\mathrm{p}}^{\mathrm{hi}*} \approx \frac{\delta_{\mathrm{P}}}{bk_{\mathrm{hi}}}, \qquad P^* \approx \frac{1}{k_{\mathrm{hi}}}\left(\gamma_{\mathrm{hi}}Q - \frac{1}{\tau}\right). \tag{35}$$

This is the fixed point mentioned in the main text. Next, we neglect the lysis and pilus expression delays to obtain an analytical estimate of the frequency of oscillations in Figure 2D. Without delays, the Equations 30 and 31 simplify to

$$\frac{\mathrm{d}B_{\mathrm{p}}^{\mathrm{hi}}(t)}{\mathrm{d}t} \approx \gamma_{\mathrm{hi}}\, QB_{\mathrm{p}}^{\mathrm{hi}} - \frac{1}{\tau}B_{\mathrm{p}}^{\mathrm{hi}} - k_{\mathrm{hi}}B_{\mathrm{p}}^{\mathrm{hi}}P, \tag{36}$$

$$\frac{\mathrm{d}P(t)}{\mathrm{d}t} \approx bkB_{\mathrm{p}}^{\mathrm{hi}}P - \delta_{\mathrm{P}}P. \tag{37}$$

The Jacobian $J$ of these simplified dynamics is

$$J = \begin{pmatrix} \gamma_{\mathrm{hi}}\, Q - \frac{1}{\tau} - k_{\mathrm{hi}}P & -k_{\mathrm{hi}}B_{\mathrm{p}}^{\mathrm{hi}} \\ bkP & bk_{\mathrm{hi}}B_{\mathrm{p}}^{\mathrm{hi}} - \delta_{\mathrm{P}} \end{pmatrix}. \tag{38}$$

Evaluated at the fixed point given by Eq. 35 above, the Jacobian is

$$J|_{P^*,\, B_{\mathrm{p}}^{\mathrm{hi}*}} = \begin{pmatrix} 0 & -\frac{\delta_{\mathrm{P}}}{b} \\ b\left(\gamma_{\mathrm{hi}}Q - \frac{1}{\tau}\right) & 0 \end{pmatrix}. \tag{39}$$

The eigenvalues of the above matrix are

$$\lambda = \pm i\sqrt{\delta_{\mathrm{P}}\left(\gamma_{\mathrm{hi}}Q - 1/\tau\right)}. \tag{40}$$

Thus, the frequency of oscillations of small amplitude around the fixed point is given by $\omega = \sqrt{\delta_{\mathrm{P}}\left(\gamma_{\mathrm{hi}}Q - 1/\tau\right)}$. Although the oscillations about the fixed point in Figure 2D are not necessarily small, using the parameters of Figure 2D ($Q = 10^8/\mathrm{ml}$, $\delta_P = 0.01/\mathrm{h}$, and $\gamma_{\mathrm{hi}} = 10^{-8}$ ml/h) and Eq. 40 predicts the oscillations to have a period $T_{\mathrm{osc}} = 2\pi/\omega \sim 6 \times 10^1$ h, which roughly agrees with the oscillation period observed in Figure 2D.

### 3 Model with heterogeneous expression of conjugation machinery

To capture heterogeneity, we describe $N = 20$ bacterial populations $B_{\mathrm{p}}^{\mu}$ (cell types $\mu \in \{1, 2, ..., N\}$) with conjugation rates $\gamma_\mu$ that are evenly spaced (on a log scale) from $\gamma_{\mathrm{lo}} = 10^{-12}$ml/h to $\gamma_{\mathrm{hi}} = 10^{-8}$ ml/h. The phage infection rate for each population is $k_\mu = \kappa\gamma_\mu$ with $\kappa = 1.0$. The dynamics include intermediate states for phage infection and pilus expression for each cell type $\mu$:

$$\frac{\mathrm{d}B_0^{\mu}}{\mathrm{d}t} = (R_0 - \delta_{\mathrm{B}})B_0^{\mu} + p_\ell R_{\mathrm{p}}B_{\mathrm{p}}^{\mu} - B_0^{\mu}\sum_{\nu=1}^{N}\gamma_\nu B_{\mathrm{p}}^{\nu} + \frac{1}{T_{\mathrm{mix}}}\left(-B_0^{\mu} + \sum_{\nu\neq\mu}\frac{B_0^{\nu}}{N-1}\right), \tag{41}$$

$$\frac{\mathrm{d}S_1^{\mu}}{\mathrm{d}t} = -\left(\frac{M}{T_{\mathrm{pili}}} + \delta_B\right)S_1^{\mu} + B_0^{\mu}\sum_{\nu=1}^{N}\gamma_\mu B_{\mathrm{p}}^{\mu}, \tag{42}$$

$$\frac{\mathrm{d}S_i^{\mu}}{\mathrm{d}t} = \frac{M}{T_{\mathrm{pili}}}S_{i-1}^{\mu} - \left(\frac{M}{T_{\mathrm{pili}}} + \delta_{\mathrm{B}}\right)S_i^{\mu}, \tag{43}$$

$$\frac{\mathrm{d}B_{\mathrm{p}}^{\mu}}{\mathrm{d}t} = \left[R_p\left(1 - p_\ell\right) - \delta_{\mathrm{B}}\right]B_{\mathrm{p}}^{\mu} + \frac{M}{T_{\mathrm{pili}}}S_M^{\mu} - k_\mu B_{\mathrm{p}}^{\mu}P + \frac{1}{T_{\mathrm{mix}}}\left(-B_{\mathrm{p}}^{\mu} + \sum_{\nu\neq\mu}\frac{B_{\mathrm{p}}^{\nu}}{N-1}\right), \tag{44}$$

$$\frac{\mathrm{d}I_1^{\mu}}{\mathrm{d}t} = Pk_\mu B_{\mathrm{p}}^{\mu} - \left(\frac{M}{T_{\mathrm{lysis}}} + \delta_{\mathrm{B}}\right)I_1^{\mu}, \tag{45}$$

$$\frac{\mathrm{d}I_i^{\mu}}{\mathrm{d}t} = \frac{M}{T_{\mathrm{lysis}}}I_{i-1}^{\mu} - \left(\frac{M}{T_{\mathrm{lysis}}} + \delta_{\mathrm{B}}\right)I_i^{\mu}, \tag{46}$$

$$\frac{\mathrm{d}P}{\mathrm{d}t} = -\delta_{\mathrm{P}} P + b\frac{M}{T_{\mathrm{lysis}}} \sum_{\mu=1}^{N} I_M^{\mu} - P \sum_{\mu=1}^{N} k_{\mu} \sum_{i=1}^{M} I_i, \tag{47}$$

$$R_0 = \alpha_0 \frac{1}{Q} \left( Q - B_0 - \sum_{\mu} B_{\mathrm{p}}^{\mu} - \sum_{i=1}^{M} \sum_{\mu=1}^{N} (S_i^{\mu} + I_i^{\mu}) \right), \tag{48}$$

$$R_{\mathrm{p}} = (1 - \Delta) R_0 \tag{49}$$

where $\mu \in \{1, 2, ...N\}$ and $i \in \{1, 2, ..., M\}$ when not summed over. With $N = 20$ and $M = 5$, this yields a total of 242 equations. The last term in Equations 40 and 42 describes the mixing of subpopulations used to regenerate the high-conjugation population after the selection event in Figure 3 of the main text. If $T_{\mathrm{mix}} \to \infty$ (as in Figure 2F of the main text), the high conjugation populations are reduced during the selection event, because they provide little benefit to the plasmid during that period (few cells available to receive plasmids), but remain under phage predation. This can result in extinction of the plasmid after the selection event ends, as shown in Figure S2.

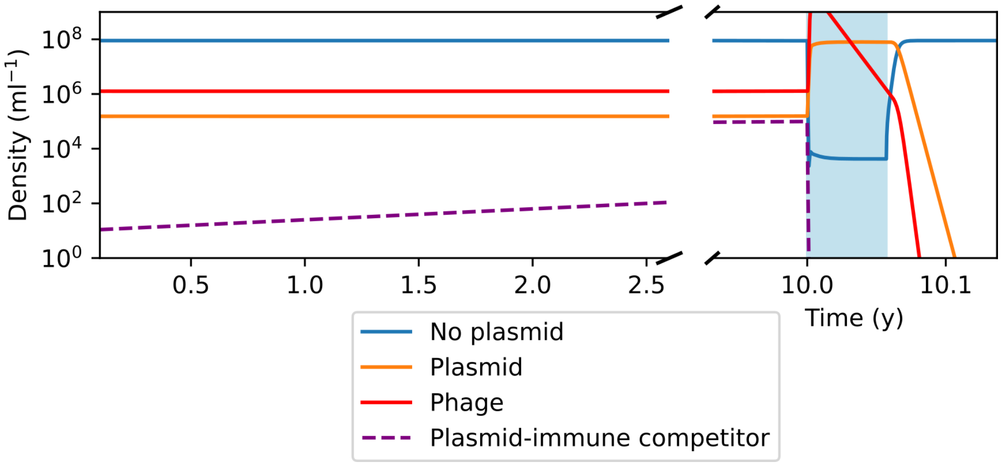

Figure S2: **Extinction of costly plasmids after selection events when high conjugation cells are not replaced.**
